# Using 2*k* + 2 bubble searches to find SNPs in *k*-mer graphs

**DOI:** 10.1101/004507

**Authors:** Reda Younsi, Dan MacLean

## Abstract

This preprint is now available in published form as: ‘Using 2*k* + 2 bubble searches to find SNPs in *k*-mer graphs’, Reda Younsi; Dan MacLean, Bioinformatics 2014; doi: 10.1093/bioinformatics/btu706

Single Nucleotide Polymorphism (SNP) discovery is an important preliminary for understanding genetic variation. With current sequencing methods we can sample genomes comprehensively. SNPs are found by aligning sequence reads against longer assembled references. De Bruijn graphs are efficient data structures that can deal with the vast amount of data from modern technologies. Recent work has shown that the topology of these graphs captures enough information to allow the detection and characterisation of genetic variants, offering an alternative to alignment-based methods. Such methods rely on depth-first walks of the graph to identify closing bifurcations. These methods are conservative or generate many false-positive results, particularly when traversing highly inter-connected (complex) regions of the graph or in regions of very high coverage.We devised an algorithm that calls SNPs in converted De Bruijn graphs by enumerating 2*k* + 2 cycles. We evaluated the accuracy of predicted SNPs by comparison with SNP lists from alignment based methods. We tested accuracy of the SNP calling using sequence data from sixteen ecotypes of *Arabidopsis thaliana* and found that accuracy was high. We found that SNP calling was even across the genome and genomic feature types. Using sequence based attributes of the graph to train a decision tree allowed us to increase accuracy of SNP calls further.Together these results indicate that our algorithm is capable of finding SNPs accurately in complex sub-graphs and potentially comprehensively from whole genome graphs.The source code for a C++ implementation of our algorithm is available under the GNU Public Licence v3 at:https://github.com/redayounsi/2kplus2

## Introduction

Single Nucleotide Polymorphisms (SNPs) between genomes of individuals are valuable markers for tracing the genetic basis of inheritable traits or diseases.Rapid detection and creation of large libraries of SNPs is vital for timely investigation and identification of genes associated with important phenotypes. Contemporary sequencing technology can sample genomes comprehensively in only hours, with these data SNP detection is typically achieved by aligning reads to a reference genome and identifying SNPs as a difference between consensus and the reference [Leggett and MacLean, 2014]. Factors such as the need for a reference sequence and the assumption of a monomorphic sample mean that the consensus approach is limited in organisms for which we lack a reference genome, in outbred diploid samples, bulked population data or analysis of metagenomes. To deal with the very large amount of sequence data that current sequencing technologies produce, De Bruijn graphs have been used to represent *k*–mer (nucleotide subsequences of arbitrary length *k*) overlap patterns in sequence reads.

These *k*–mer graphs have been implemented into efficient data structures for large collections of *k*–mers and have proven to be of great utility as the underlying data model over which numerous *de novo* genome assembly algorithms have been implemented ([Pevzner *et al.*, 2001], [Zerbino and Birney, 2008], [Simpson *et al.*, 2009]). Methods based on the De Bruijn graph have been implemented recently that can efficiently identify sample differences in sequence read sets ([Iqbal *et al.*, 2012], Leggett *et al.*, 2013]). Iqbal et al. ([Iqbal *et al.*, 2012]) produced the first model and *de novo* assembly algorithms for variant discovery and genotyping directly from sequence data without using a reference genome. They incorporated a colour attribute for the edges in the graph that represents the sample from which the sequence is derived. In these multi-sample graphs polymorphisms appear as bubbles, a closing bifurcation of length 2*k* + 2 with separate colours on each branch of the bifurcation 1. Discovering bubbles is a promising tactic for identifying SNPs in the graph, the bubble caller in Iqbal *et al* and the more computationally intensive depth-first search method in Leggett et *al.* ([Leggett *et al.*, 2013]) proceeds by marking the vertices in the graph that have at least two edges departing from them as starting points and following vertices sequentially until a vertex with at least two edges entering it is reached (see Figure 1). An alternative microassembly approach ([Peterlongo *et al.*, 2010]) begins by producing a tree of *k*–mers for an input read set picking a seed *k*–mer and assuming that it lies on one path through a SNP and then looks for an opposite *k*–mer, one substitution different, which would lie on another path through the bubble. If this can be found in the *k*–mer tree, then a recursive algorithm builds paths left and right of each *k*–mer until they join or no *k*–mer can be found. Further to graph structure, the attributes of the sample and sampled sequence reads can be used

**Figure 1:**
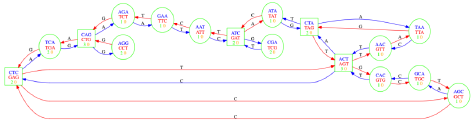
De Bruijn graph fragment with edge and vertex attributes as represented in the Cortex software and used in this study. Vertices represent *k*–mers from sequence fragments and their reverse complements, edges represent directional overlap between sequence and edge annotations represent the changed base between edges. Coloured numbers in vertices indicate coverage (times the *k*–mer is observed) in the respective ‘orange’ or ‘green’ coloured sample.

## 2 Approach

De Bruijn graphs of sequence reads are typically very large with the vast majority of vertices having only a single in and out edge, as such no complex bubbles can exist in these areas and much of the graph need not be searched. Thus, we first build a sub-graph from a branching vertex that has at least two out edges and search sub-graphs for cycles. We retain only cycles of length 2*k* + 2 with two equidistant branching vertices in which the edge paths on each branch are a different colour. These constraints allows us to simplify the search for cycles by only checking the locality of the starting branching vertex. Once the algorithm terminates, a simple walk of the vertices in the bubble, collecting the labels, gives the sequence. The two coloured *k*+1 walks result in separate nucleotide sequences of length *k* + 1 in which the first nucleotide differs between the samples. Our method’s very precise description of a bubble means that it has the potential to be extremely specific and generate highly accurate lists of SNPs from graph structure alone.

## 3 Methods

### 3.1 Data sets

We used sixteen sets of Illumina sequence reads from different ecotypes of the model plant *Arabidopsis thaliana* with the *Arabidopsis thaliana* ecotype Col-0 genome as the common reference ([AGI, 2000]). For each of these high-quality SNP lists are available ([Xiangchao *et al.*, 2011]). *Arabidopsis thaliana* has a small (126Mbp) tractable genome with only a relatively small repeat content and well catalogued genetic diversity, making it an ideal test organism.

### 3.2 2*k* + 2 Bubble detection algorithm

We used Cortex [Iqbal *et al.*, 2012] to build a graph directly from sequence data, removing paths of vertices with coverage 2 or below and tips less than 100 nucleotides in length. The resulting graph is exported and is then transformed to an undirected graph, retaining the edge attributes. Then 2*k* + 2 (Algorithm 1) is applied to create short contig bearing SNPs.

### 3.3 Using canonical SNP lists to assess accuracy of the algorithm

In order to assess the accuracy of the SNPs predicted by 2*k* + 2 we compared our predictions against published lists of SNPs called in the [?] study. The contigs generated by the 2*k* + 2 algorithm were used as query sequence in a BLASTn search with default settings [Altschul *et al.*, 2000] against the *A.thaliana* Col–0 reference sequence and used the top hit in the BLASTn to find the position of the predicted SNP in the genome. By comparing the position of the published SNPs with those from our algorithm we could then calculate the number of SNPs accurately predicted by our algorithm as the number found in the published lists. We used three measures of accuracy; sensitivity (True positive / True Positive + False Negative), specificity (True negative / True negative + False Positive) and accuracy (proportion of predicted SNPs that were included in the known SNP list). Specificity in this analysis is always very close to 100 % because of the very large number of non-SNP genome sites.

**Algorithm 1.**
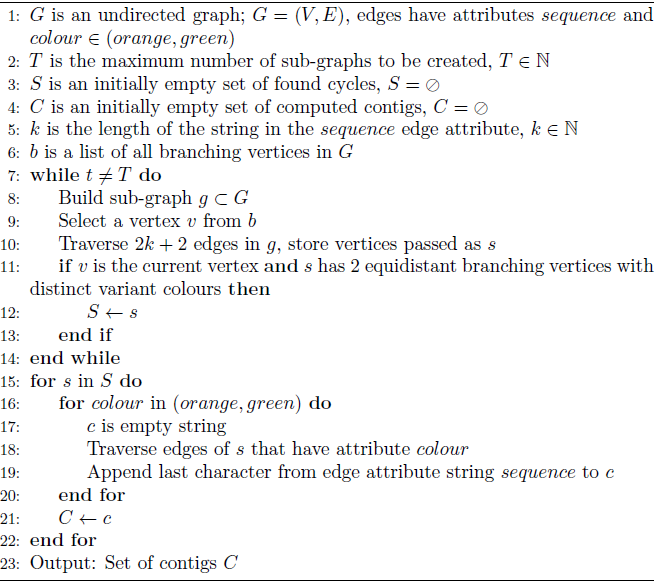
A bubble detection algorithm (2*k* + 2).

### 3.4 Using bubble attributes to filter and improve accuracy

In order to use a classification algorithm such as decision trees [Quinlan, 1986], 2*k* + 2 bubble edge attributes were used to create a vector of values describing each bubble. We used coverage, summed coverage over each branch of the bubble, mean coverage of the bubble and the mean number of branching vertices of the sub-graph where the bubble resides (See Supplemental Table ?? for summaries of the datasets used for classification). Each bubble in a graph was classified as a Real SNP or a False Positive according to its presence in the SNP list and were divided into a training set and testing set. 2/3 of the dataset is used for training the classifier and 1/3 for testing. To train we used a decision tree found in the freely available WEKA package [Mark *et al.*, 2009] and kept the default parameters of the classifier.

## 4 Results and discussion

### 4.1 SNP prediction based on structure alone is highly accurate

We ran the 2*k* + 2 algorithm on graphs built with *k* = 21 and predicted SNPs in the sixteen *A. thaliana* ecotypes. On average we predicted 146741.9 ± 18183.2 SNPs. The total number of SNPs we predicted increases fairly linearly with the genetic distance of the test ecotype from the reference Col–0 (using the number of canonical SNPs between Col–0 and each ecotype as a proxy for distance, Fig. 2B) and we achieved mean accuracy of 83.7% ± 3.5% of correctly predicted SNPs (Fig. 2B), indicating that bubble retrieval with our bubble search algorithm alone produces highly accurate SNP predictions and that the algorithm can detect more SNPs when there are more to be detected. The genetic distance of the ecotype from the Col–0 reference does not have a marked effect on accuracy in the majority of cases we examined. For 12 of the ecotypes we observed accuracy clustered around the 85% mark (Fig. 2C), indicating a natural limit on the accuracy of the prediction when using bubbles. Four of the ecotypes had lower than 80% accuracy.

**Figure 2:**
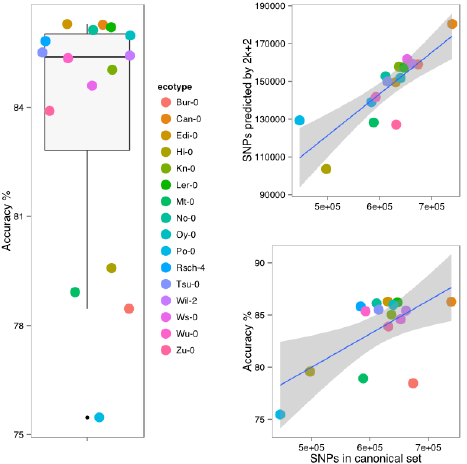
Accuracy of SNP prediction using the 2*k* + 2 algorithm on 16 A. *thalian* ecotypes. Panel A (left side) shows distribution of accuracy over all 16 ecotypes. Panel B (top right) gives the number of SNPs predicted found by 2*k* + 2 as a function of the SNPs expected from the canonical list. Panel C (bottom right) gives the number of SNPs accurately predicted as a function of the number of SNPs in the canonical set.

### 4.2 Accuracy of snp calling is consistent across the genome

To determine whether our strategy suffered from bias toward particular genomic regions, we compared the position of SNPs predicted by the 2*k* + 2 structure with those from the canonical lists for Col–0 and Tsu–1. Analysis of the proportion of published SNPs found by 2*k* + 2 in windows of 20 kbp across the genome showed consistently high recovery of SNPs. Average recall in the windows was 73%. (Fig. 3). The number of published SNPs detected fell substantially in regions corresponding to the centromeres.

**Figure 3:**
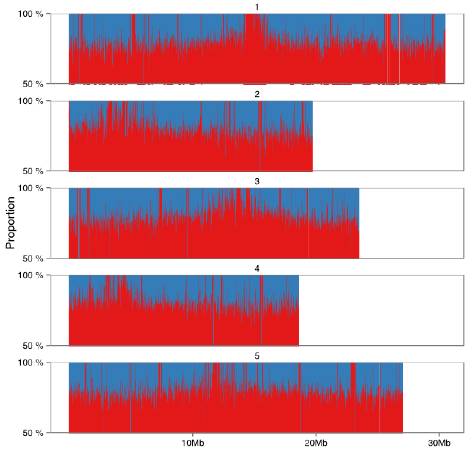
The proportion of consensus-called SNPs predicted by the *2k* + 2 algorithm in 20 kb windows of the five *Arabidopsis thaliana* nuclear chromosomes SNPs were called on graphs composed of reads from ecotype Tsu-1 relative to the Col-0 common reference. Red peaks indicate the proportion of all SNPs (blue area) found by 2k + 2 in each window.

To determine whether SNPs in any particular genomic feature types were preferentially missed by our algorithm we examined enrichment of feature types in the 2.5% least sensitive 20 kbp windows. The distribution of sensitivity estimates in the windows was observed to be approximately normal and we applied a Bonferroni-corrected hypergeometric test to each feature type, comparing the proportion in the TAIR 9 Gene Ontology annotation of the genome with the proportion in the bottom 2.5% of sensitivity windows. We saw that non-coding regions such as pseudogenes and transposable elements were enriched in the sample (*p* < 0.001) (Table 1) it is likely because centromeric regions are enriched in these features ([AGI, 2000]). The lack of bias toward successful SNP calls in any particular genomic feature or region indicates that the random search strategy we are using to create sub-graphs does not result in bias toward any specific parts of the genome and we conclude that the 2*k* + 2 algorithm has the potential to be part of a general SNP finding pipeline.

### 4.3 The 2*k* + 2 algorithm finds SNPs in complex portions of the graph

To establish the performance of the algorithm in complex regions of the graph we compared the total number of branching vertices in a 100 vertex sub-graph with the number of SNPs that were predicted when each sub-graph was searched exhaustively. We looked at 54,616, 35,043 and 36,483 randomly sampled subgraphs with more than two branching vertices for ecotypes Can-0, Bur-0 and Po-0 respectively. The majority of sub-graphs have just a few branching vertices though there are a few very heavily branched sub-graphs with more than 20 out of 100 vertices with branches (Figure 4A). The number of SNPs predicted in each sub-graph is most usually 1 (49 %, 36 % and 43 % of subgraphs contain only 1 SNP for Can–0, Bur–0 and Po–0, see supplemental data) regardless of the number of branching vertices. Intuitively we would expect the total number of SNPs predicted to increase as complexity increases as SNPs nearby to each in the genome other would increase graph complexity but this is not observed and for sub-graphs with more than 15 branching vertices we predict only one SNP in any ecotype 4. Above 90 % of predicted SNPs are in sub-graphs with only 2–5 branching vertices Figure 4B and supplemental data). The result indicates that the algorithm is most successful at retrieving SNPs in sub-graphs with only two branching vertices, though can handle moderately complex graphs and successfully identify SNPs within those sub-graphs. We have not been able to confirm whether SNP density does truly increase with graph complexity and therefore cannot rule out whether SNP prediction drop off in more complex regions is a true property of the graph or a failure of the*2k* + 2 algorithm to detect in these regions.

**Figure 4:**
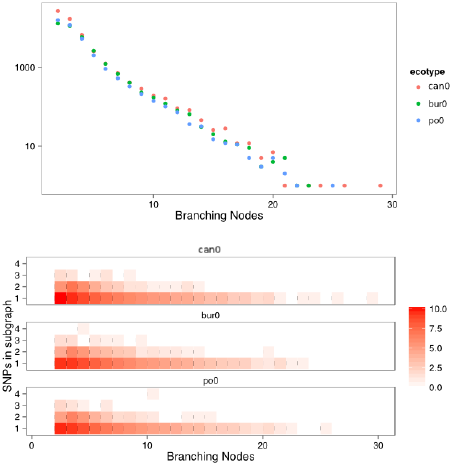
Distribution of number of subgraphs with given number of branching nodes (A, top panel) and the number of SNPs predicted in all sub-graphs with given number of branching nodes (B, bottom panel). Data from three ecotypes Can-0, Bur-0 and Po-0 are presented. Colour scale represents natural log of counts, see supplemental data for real values.

**Table I.** Genomic feature enrichment in the *Arabidopsis thaliana* genomic 20 kb windows that had the lowest 2.5% called SNP sensitivity rates in Col–0/Tsu–1 data. For each feature type the proportion of all features in the genome and in the sample and a Bonferroni corrected *p*-value from the hypergeometric test is presented.

| Feature | Genome Proportion | Sample Proportion | $p$ |
| --- | --- | --- | --- |
| Exon | 0.74 | 0.72 | 1 |
| 5' UTR | 0.12 | 0.08 | 1 |
| 3' UTR | 0.11 | 0.11 | 1 |
| miRNA | 0.00 | 0.00 | 1 |
| tRNA | 0.00 | 0.00 | 1 |
| ncRNA | 0.00 | 0.01 | <0.001 |
| Pseudogene | 0.00 | 0.01 | <0.001 |
| Pseudogenic transcript | 0.00 | 0.01 | <0.001 |
| Pseudogenic exon | 0.00 | 0.01 | <0.001 |
| Transposable element gene | 0.01 | 0.05 | <0.001 |
| snoRNA | 0.00 | 0.00 | 1 |
| snRNA | 0.00 | 0.00 | 1 |
| rRNA | 0.00 | 0.00 | 1 |

### 4.4 Accuracy is improved by filtering candidate bubbles using a decision tree

We trained a decision tree to distinguish between known SNPs and non-SNP bubbles and applied it to the predicted bubbles from 16 ecotypes to successfully classify bubbles that represented real SNPs with higher accuracy whilst maintaining similar sensitivity and specificity than using bubble structure alone (Figure 5). The specificity ranged between 97.22 % and 86.33 % and sensitivities vary between 84.63 % and 68.71 % each of which is similar to the values seen when using structure alone. Accuracy in the predicted SNP set was from 86.33 % to 97.04 %, much higher than from structure. This result indicates that structure and extra bubble attributes can give very highly accurate sets of predicted SNP sets. In a similar approach, with very similar A. *thaliana* data Leggett *et al* ([Leggett *et al.*, 2013]) used a ranking heuristic to classify SNPs after initial prediction but found that accuracy of that heuristic was compromised as the number of SNPs predicted increased and in SNP sets of over 100000 each newly predicted SNP has only a 40 % likelihood of being accurate such that SNP sets of the size we predict contained very high numbers of false positive SNP calls, the 2*k* + 2 algorithm and classifier outperforms that approach significantly returning better than 90 % accurate SNP calls for very large SNP sets.

**Figure 5:**
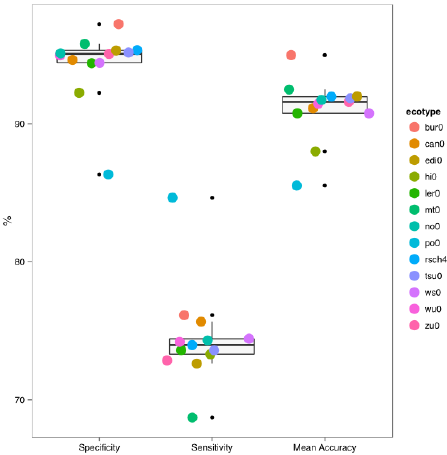
Classification accuracy of SNPs called by using the Decision Tree algorithm to classify bubbles found in 2*k* + 2 searches on graphs made from sequence data of 13 ecotypes of *Arabidopsis thaliana.* The results are the mean of 10 runs using the decision tree algorithm in WEKA [Mark *et al.*, 2009]. Results are the mean of correctly classified bubbles that contain SNPs as listed in [Xiangchao *et al.*, 2011].

## 5 Conclusion

The 2*k* + 2 algorithm takes a graph theoretical approach to identifying topological structures, namely *2k* + 2 cycles in undirected graphs that can represent SNPs in sequences input to a donor De Bruijn graph. We have shown that 2*k* + 2 can be used to generate sets of SNPs in graphs at high accuracy in the *Arabidopsis thaliana* genome which is increased by application of the decision tree classifier. 2*k* + 2 found SNPs across all portions of the *Arabidopsis thaliana,* genome without bias toward feature type or region. We did not find that all SNPs seen in alignment based methods using a reference could be detected with this algorithm and in all our experiments we found that around 70 % of SNPs seen by alignment could be predicted. The 2*k* + 2 strategy is therefore a useful algorithm that should find general use in reference-free SNP calling strategies in the future.

## Acknowledgements

We wish to thank Ricardo Ramirez-Gonzalez and Dr Graham Etherington for assistance during this project.

Funding:RY was funded by a BBSRC (Biotechnology and Biological Sciences Research Council) TRDF2 grant (ref: BB/I023798/1) to DM. DM was supported by The Gatsby Charitable Foundation.

